# Serine-arginine protein kinase 1 regulates Ebola virus transcription

**DOI:** 10.1101/717371

**Authors:** Yuki Takamatsu, Verena Krähling, Larissa Kolesnikova, Sandro Halwe, Clemens Lier, Stefan Baumeister, Takeshi Noda, Nadine Biedenkopf, Stephan Becker

## Abstract

Ebola virus (EBOV) causes a severe and often fatal disease for which no approved vaccines or antivirals are currently available. EBOV transcription requires the sequential phosphorylation and dephosphorylation of the viral transcription factor VP30. While dephosphorylation is carried out by phosphatases PP2A and PP1, the VP30-specific kinase is unknown. Here, we report that serine-arginine protein kinase 1 and 2 (SRPK1 and SRPK2) phosphorylate serine-29 of VP30, which is located in an N-terminal R_26_xxS_29_ motif. Interaction with VP30 via the R_26_xxS_29_ motif recruits SRPK1 into EBOV-induced inclusion bodies, the sites of viral RNA synthesis and an inhibitor of SRPK1/SRPK2 downregulates primary viral transcription. When the SRPK1 recognition motif of VP30 was mutated in a recombinant EBOV, virus replication was severely impaired. It is presumed that the interplay between SRPK1 and PP2A in the EBOV inclusions provides a comprehensive regulatory circuit to ensure the activity of VP30 in EBOV transcription.

## Introduction

Reversible phosphorylation is an important posttranslational process that regulates protein conformation, thereby promoting protein-protein interactions, signal transduction and regulating protein synthesis and degradation (1-3). Similarly, for many important human RNA viruses, phosphorylation of viral proteins is invoked to regulate transcription, replication, and virus assembly (4-10).

Ebola virus (EBOV), a filovirus, causes severe hemorrhagic fever with high fatality rates and is mainly treated symptomatically due to the lack of approved antivirals. Nonetheless, promising EBOV inhibitors are currently at different stages of clinical development (11). In addition to antivirals that directly target viral enzymes or surface proteins, the development of inhibitors against proviral host cell factors has become an attractive option because it is suspected that such inhibitors are less prone to inducing resistance (12). However, to develop such host-targeting antivirals, it is necessary to understand the molecular mechanisms at the host/virus interface.

EBOV entry is accomplished by macropinocytosis, resulting in the release of viral nucleocapsids from the endosomal/lysosomal compartment into the cytoplasm (13). The nucleocapsid then acts as a template for primary transcription, which is executed by nucleocapsid-associated polymerase L, the polymerase cofactor VP35 and the transcription factor VP30 (14, 15). After viral mRNAs have been translated and sufficient amounts of viral proteins have been produced, replication and encapsidation of the genomic RNA occurs. At the same time, secondary transcription takes place, which is distinct from primary transcription by the abundance of viral proteins. Although the precise mechanism by which VP30 supports primary and secondary transcription is unknown, it is clear that this function is regulated by the phosphorylation status of VP30 (10, 16). VP30 phosphorylation enhances its binding to the nucleoprotein NP while inhibiting its interaction with RNA and VP35, collectively resulting in downregulation of viral transcription (10, 16-18). Interestingly, a nonphosphorylatable mutant of VP30 is sufficient to support secondary transcription, whereas primary transcription requires sequential phosphorylation and dephosphorylation (19). Six N-proximal serine residues of VP30 are potential phosphoacceptor sites. Intriguingly, a mutant of VP30 (VP30^29S^) in which five of the six serine residues are replaced by alanine, with only serine-29 remaining, was found to be capable of supporting both primary and secondary transcription. Indeed, recombinant EBOV (recEBOV_29S) expressing VP30^29S^ instead of VP30^wt^ was rescuable and behaved in a manner similar to wild-type EBOV (19), suggesting that reversible phosphorylation of serine-29 was necessary and sufficient to support all functions of VP30 during transcription.

Previous work has shown that the cellular phosphatases PP1 and PP2A are able to dephosphorylate VP30 (10, 18, 20). Dephosphorylation of VP30 by PP2A was found to be mediated by NP, which simultaneously recruited VP30 and PP2A into viral inclusion bodies via two adjacent binding motifs. The close proximity of VP30 and PP2A in association with NP results in efficient VP30 dephosphorylation and consequent activation of EBOV transcription (21, 22).

Although VP30’s site-specific dephosphorylation has been well elucidated, the nature of VP30-specific kinase(s) and how and where VP30 phosphorylation occurs remain completely unknown. Using a proteomics approach, we identified two host kinases, serine-arginine protein kinase 1 and 2 (SRPK1 and SRPK2), that specifically phosphorylate the important residue serine-29 of VP30. Ectopic expression of SRPK1 enhanced VP30 phosphorylation and thus downregulated EBOV transcription, therefore reducing the propagation of infectious EBOV. Inhibition of endogenous SRPK1 downregulated primary transcription. We further identified the R_26_xxS_29_ motif in VP30 as a main SRPK1 recognition motif and confirmed the importance of this motif for viral growth. Together with the previously detected VP30-specific cellular phosphatase PP2A, the newly described VP30-specific kinase SRPK1 represents a phosphorylation/dephosphorylation circuit that regulates EBOV mRNA synthesis in viral inclusion bodies.

## Results

### Identification of a VP30-specific cellular kinase

VP30 is phosphorylated at six N-proximal serine residues (Figure 1A) (10); the most important is serine-29, the phosphorylation and dephosphorylation of which are necessary to ensure VP30’s role in primary and secondary transcription (19).

**Figure 1.**
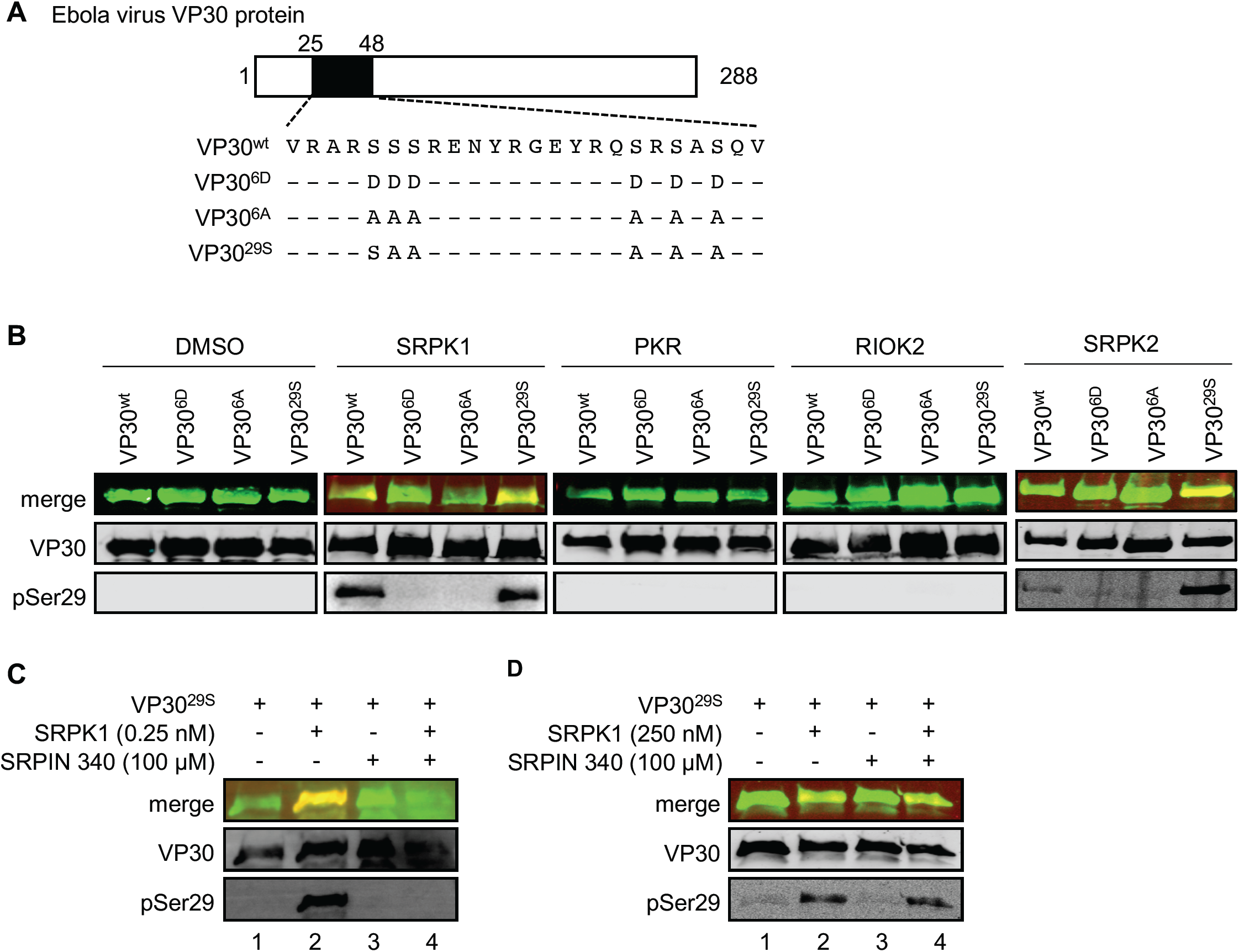
Identification of VP30-specific kinases. (A Schematic drawing of Ebola virus VP30 showing amino acid sequence 25-48 of VP30 and the mutants of VP30 employed. (B, C) Bacterially expressed MBP fusion proteins of VP30 (amino acids 8-272) or phosphomimetic mutants of VP30 were phosphorylated *in vitro* using recombinant SRPK1, PKR, RIOK2, and SRPK2. Samples were subjected to SDS-PAGE and western blotting. VP30 and phosphorylated VP30 (serine 29) were detected using a guinea pig anti-VP30 and a rabbit anti-pSer29 antibody. (B) Bacterially expressed VP30 and mutants of VP30 were incubated with either DMSO (control), recombinant SRPK1, recombinant PKR, recombinant RIOK2, or recombinant SRPK2 in kinase reaction buffer for 30 min at room temperature. (C) Bacterially expressed VP30^29S^ was incubated with either SRPK1 (0.25 nM), SRPIN340 (100 µM), or a mixture of SRPK1 (0.25 nM) and SRPIN340 (100 µM) in kinase reaction buffer for 18 h at room temperature. (D) Experimental setting as in (c), except the amount of recombinant SRPK1 was increased to 250 nM.

To identify EBOV VP30-associated kinase(s), we adopted a proteomics approach by employing different versions of VP30: wild-type VP30 (VP30^wt^); a nonphosphorylatable mutant of VP30 in which all six major phosphorylation sites at the N-terminus are mutated to alanine (VP30^6A^); a mutant that mimics fully phosphorylated VP30 via replacement of the six serines to negatively charged aspartic acid residues (VP30^6D^); and VP30^29S^ (Figure 1A) (10, 19). Ectopically expressed FLAG-tagged VP30 mutants were immunoprecipitated, and coprecipitating cellular proteins were eluted (16) and digested with trypsin prior to analysis by liquid chromatography-tandem mass spectrometry (Supplementary Figure 1) (23). A number of kinases coprecipitated with VP30 (Figure 1B, Figure supplement 1); of these, we focused on SRPK1, interferon-induced double-stranded RNA-activated protein kinase (PKR) and serine/threonine-protein kinase RIO2 (RIOK2). The three kinases displayed high binding with VP30^wt^ and VP30^29S^, with lower binding to VP30^6A^ or the FLAG-epitope alone (Figure 1B, Figure supplement 1). We then performed *in vitro* phosphorylation assays to examine whether the kinases are able to phosphorylate VP30. Bacterially expressed and purified VP30 mutants were incubated with the respective recombinant kinases in the presence of ATP, and VP30 phosphorylation was monitored using an anti-pSer29 antibody that specifically recognizes phosphorylated serine at position 29 (20). SRPK1 and the highly related SRPK2 clearly phosphorylated VP30^wt^ and VP30^29S^, whereas PKR and RIOK2 did not (Figure 1b). To confirm that phosphorylation of VP30 was specifically executed by SRPK1, we applied the SRPK1/SRPK2-specific inhibitor SRPIN340 (24), which resulted in a significant decrease in phospho-VP30^29S^ (Figure 1C, lanes 3, 4). Moreover, the inhibitory effect of SRPIN340 was dependent on the ratio to SRPK1, as increasing the amount of SRPK1 outcompeted the influence of SRPIN340 (Figure 1D, lane 4). Altogether, mass spectrometry analyses, *in vitro* phosphorylation assays and a specific inhibitor identified SRPK1/SRPK2 as VP30-specific kinases able to phosphorylate the important serine at position 29 in VP30.

We then analyzed intracellular phosphorylation of VP30 and found that VP30 was maintained in a nonphosphorylated state in the absence of phosphatase inhibitors (Figure 2A, lanes 1 – 5). Indeed, phosphorylated VP30^wt^ and VP30^29S^ was only observed after treatment of VP30-expressing cells with okadaic acid (OA), an inhibitor of VP30-specific phosphatases PP1 and PP2A (Figure 2A, lanes 7, 10). Similarly, phosphorylation of VP30^wt^ and VP30^29S^ was detected upon ectopic expression of SRPK1, suggesting that the phosphatase activity acting on VP30 was outcompeted by SRPK1 overexpression and that the phosphorylation state of VP30 was shifted toward the phosphorylated form (Figure 2A, lanes 12 and 15). Furthermore, phosphorylation of VP30 strongly increased when PP2A was inhibited by OA and SRPK1 was ectopically expressed (Figure 2B, lanes 1 and 2), and VP30 phosphorylation was inhibited in the presence of OA and SRPIN340, indicating that SRPK1 is a major kinase that phosphorylates VP30 at position serine-29 (Figure 2B, lanes 2 and 3). As SRPIN340 treatment partially inhibited VP30 phosphorylation in the presence of ectopically expressed SRPK1, it appears that although the chosen concentration of SRPIN340 was sufficient to efficiently counteract endogenous SRPK1, it was not able to completely inhibit ectopically expressed SRPK1 (Figure 2B, lanes 2 and 4). Quantification of three independent experiments confirmed the statistical significance of the observations (Figure 2B, graph). In summary, SRPK1 specifically phosphorylates VP30^wt^ and VP30^29S^ *in vitro* and in cultured cells and regulates the phosphorylation state of VP30 together with OA-sensitive phosphatases, likely PP2A and/or PP1.

**Figure 2.**
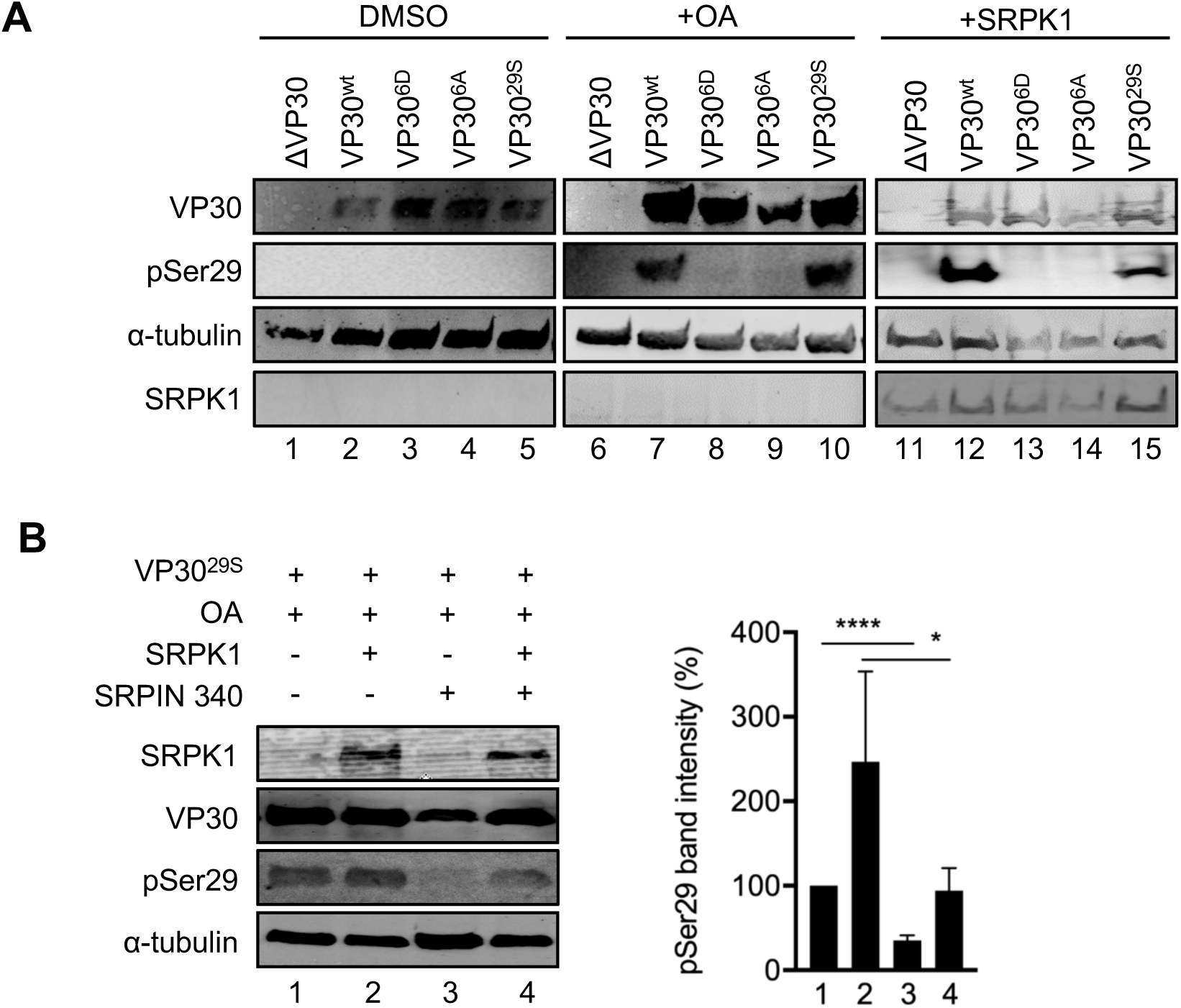
SRPK1 phosphorylation of VP30 in cells. (A, B) HEK293 cells expressing VP30 or mutants of VP30 were treated with either OA or transfected with a plasmid encoding SRPK1 and lysed at 24 h p.t. Cell lysates were subjected to SDS-PAGE and western blotting using antibodies against VP30, pSer29, α tubulin and SRPK1. (A) Lanes 1-5: DMSO-treated cells (ctrl) incubated for 18 h. Lanes 6-10: Cells incubated with 25 nM OA (inhibitor of PP1/PP2A) for 18 h. Lanes 11-15: VP30-expressing cells were transfected with 500 ng of a plasmid encoding SRPK1. (B) Phosphorylation of VP30^29S^. HEK293 cells expressing VP30^29S^ were treated with 25 nM OA and DMSO (control, lane 1), 500 ng SRPK1 (lane 2), 30 µM SRPIN340 (lane 3), or both 500 ng SRPK1 and 30 µM SRPIN340 (lane 4). Quantification of the pSer29 band intensity in lanes 1-4 is shown in the graph. The pSer29 signal was normalized to the VP30 signal. The band intensity of the control (lane 1) was set as 100%. Numbers represent the mean and SD of three independent experiments. Asterisks indicate statistical significance; * p < 0.05, **** p < 0.0001.

### SRPK1 regulates EBOV genome transcription/replication

We next used an EBOV-specific minigenome assay to analyze whether SRPK1 influences the transcriptional support activity of VP30 (14). The activity of SRPK1 was expected to promote the presence of phosphorylated VP30, which does not support transcription (10). Indeed, although ectopic expression of SRPK1 had no effect on reporter gene levels in the presence of a nonphosphorylatable VP30 (VP30^6A^), SRPK1 decreased reporter activities in the presence of VP30^wt^ or VP30^29S^ in a dose-dependent manner (Figure 3a). As a negative control, we used the phosphomimetic mutant VP30^6D^, which is unable to support viral transcription (10).

**Figure 3.**
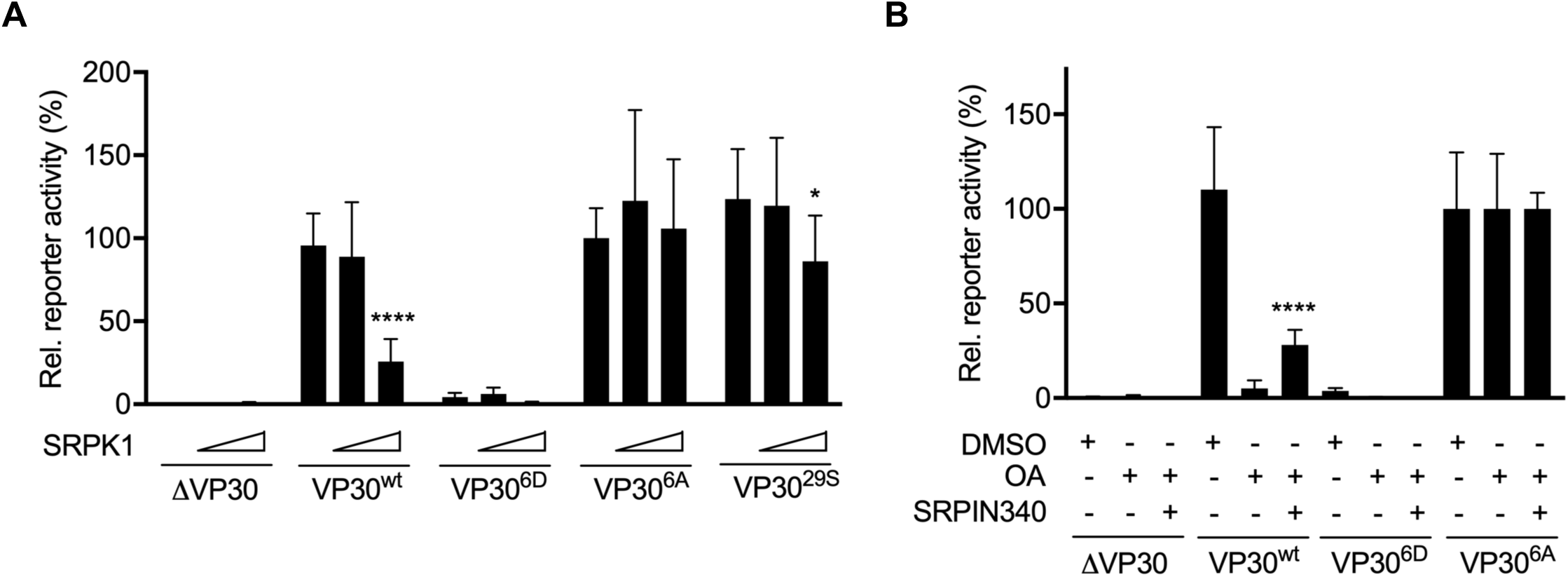
SRPK1 regulation of EBOV transcription/replication. (A) Dose-dependent effect of SRPK1 in the EBOV-minigenome assay. HEK293 cells were transfected with plasmids encoding EBOV minigenome assay components (NP, VP35, L, the EBOV-specific minigenome, T7 polymerase, absence or presence of each VP30 phenotype, and a plasmid encoding firefly luciferase for normalization). Different amounts of the SRPK1-encoding plasmid (0, 100, or 500 ng) were cotransfected. The results obtained with the phosphorylation-independent VP30^6A^ were set to 100%. Samples prepared in the absence of VP30 (ΔVP30) represented the baseline levels of the assay. (B) Effects of specific inhibition of SRPK1 by the inhibitor SRPIN340 on EBOV transcription/replication. HEK293 cells were transfected with plasmids encoding EBOV minigenome assay components (see above). At 30 h p.t., cells were incubated with either DMSO (control), 25 nM OA, or a mixture of 25 nM OA and 30 μM SRPIN340. At 48 h p.t., cells were lysed, and reporter gene activity was analyzed as indicated in (A). The mean and SD from three independent experiments are indicated. Asterisks indicate statistical significance; *p < 0.05, ****p < 0.0001.

To monitor the effect of endogenous SRPK1, we used OA to inhibit the highly active VP30-specific phosphatases PP2A and PP1, which resulted in hyperphosphorylated VP30 (Figure 2A) and consequently decreased VP30^wt^-mediated reporter gene activity by more than 90% (Figure 3B, VP30^wt^, middle bar) (10, 19). Treatment of cells with both, OA and SRPIN340 partially restored reporter gene activity (Figure 3b, VP30^wt^; right bar), indicating that SRPK1 and PP2A and/or PP1 together target VP30 and determine VP30’s phosphorylation status (as shown in Figure 2B) and, consequently, regulate viral transcription (19, 20). In the absence of OA and thus the presence of VP30-specific phosphatase activity, inhibition of endogenous SRPK1 by SRPIN340 had no effect on secondary viral transcription/replication (Supplementary Figure 2). This result was consistent with previous reports demonstrating that VP30 is efficiently dephosphorylated by PP2A in the presence of NP, resulting in a highly active VP30 and efficient secondary viral transcription in a minigenome assay (20, 22). Therefore, it was not surprising that the already dephosphorylated VP30 cannot be further activated by the inhibition of endogenous SRPK1. To evaluate the cytotoxicity of SRPIN340, we performed cell viability assays in the presence of increasing concentrations of the inhibitor (Supplementary Figure 2). Cytotoxicity was negligible at the selected concentration (30 µM), as previously reported (24, 25).

Together, these findings indicate that SRPK1 plays an important role in EBOV transcription/replication by modulating the VP30 phosphorylation status.

### The role of VP30 phosphorylation in primary transcription

To investigate the functional significance of VP30 phosphorylation by endogenous SRPK1/SRPK2 for primary transcription, we used an EBOV-specific transcription- and replication-competent virus-like particle assay, trVLP assay. All EBOV proteins and an EBOV-specific minigenome were ectopically expressed, resulting in the formation of nucleocapsid-like structures that were transported to the plasma membrane and released into the supernatant in the form of trVLPs. The released trVLPs were used to infect fresh target cells (indicator cells) to monitor primary viral transcription (Figure 4A) (26-28). Here, we compared SRPIN340-treated with mock-treated indicator cells. Interestingly, treatment with SRPIN340 decreased the reporter gene activities for trVLPs that contained VP30^wt^ to 78%, and for trVLPs that contained VP30^29S^ to 53% (Figure 4B). Though inhibitory activity of SRPIN340 was statistically significant, primary transcription was not completely blocked suggesting the existence of additional VP30-specific kinases. This result is consistent with an important role of SRPK1 or SRPK2 for efficient primary transcription of EBOV (19).

**Figure 4.**
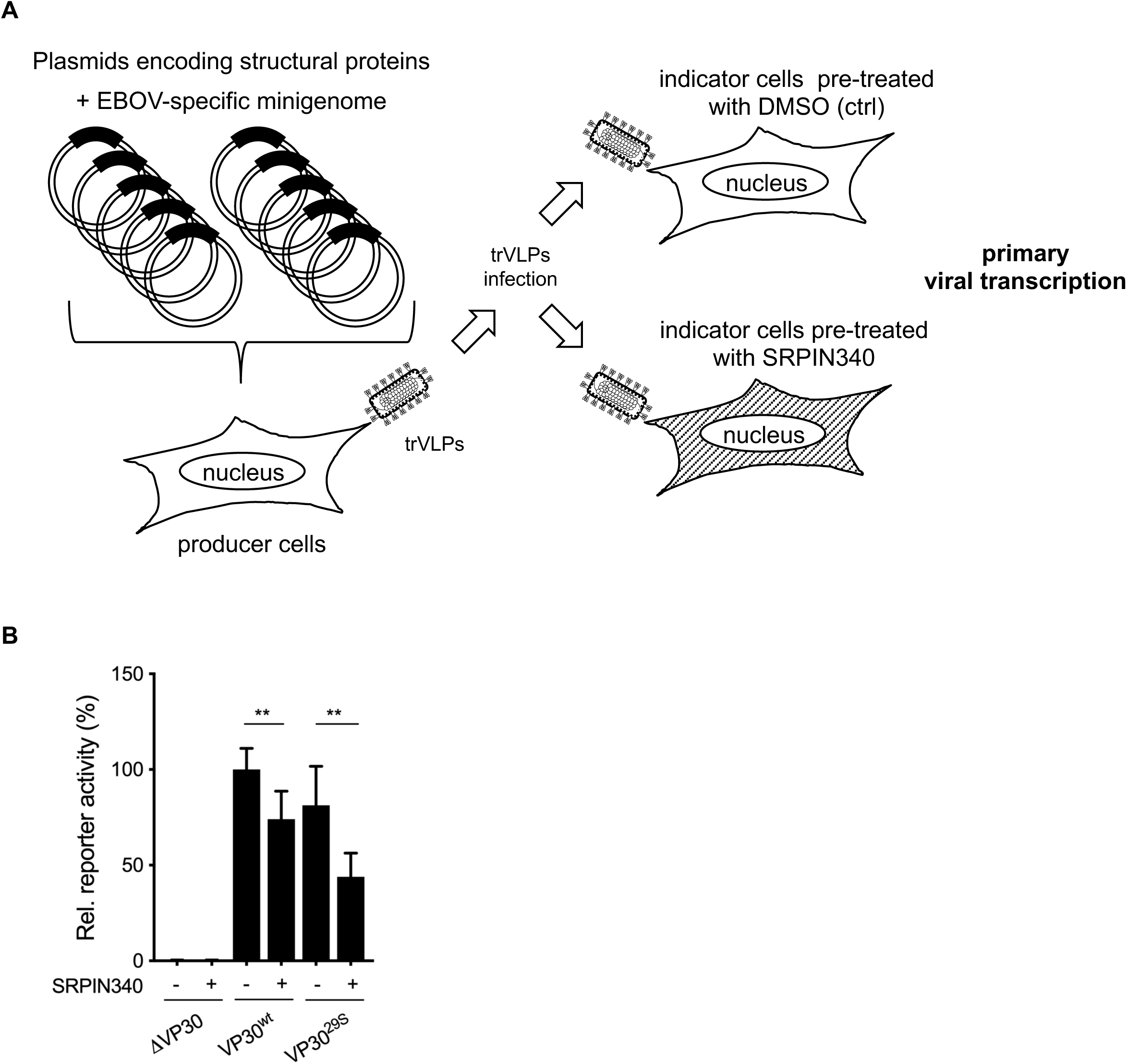
The role of SRPK1 -mediated VP30 phosphorylation in primary transcription. (A) A schematic of EBOV transcription- and replication-competent virus-like particle (trVLP) assay. HEK293 cells (producer cells) were transfected with plasmids expressing all of the viral structural proteins and an EBOV-specific minigenome encoding Renilla luciferase together with plasmid encoding firefly luciferase for normalization. The replicated minigenomes are released in the form of virus-like particles (trVLPs) into the supernatant, which was collected and purified via sucrose cushion at 72 h p.t. Huh-7 cells, the indicator cells, were pre-treated either with DMSO (ctrl) or 30 µM SRPIN340 for 24 hours. The purified trVLPs were used to infect pre-treated Huh7 cells, and reporter gene activities were measured at 48 h p.t. (B) Relative reporter activities in indicator cells at 48 h p.i. The luciferase activity of VP30^wt^-expressing cells treated with DMSO was set to 100%. Samples prepared in the absence of VP30 (ΔVP30) represented the baseline levels of the assay. The mean and SD from three independent experiments are indicated. Asterisks indicate statistical significance; *p < 0.05, ***p < 0.001.

### SRPK1 activity regulates EBOV infection

Subsequently, we analyzed the effect of ectopically expressed SRPK1 on EBOV infection using recombinant EBOVs expressing either VP30wt (recEBOV_wt) or VP30^29S^ (recEBOV_29S (19)). Plasmids encoding SRPK1 were transfected into HEK293 or HUH7 cells 12 h prior to EBOV infection and cell supernatants were collected at one and two days postinfection (p.i.). The titer of infectious EBOV in the supernatants was determined using a TCID50 assay (29). Ectopic expression of SRPK1 in HEK293 cells and Huh-7 cells resulted in a reduction in the virus titer by approximately one log (Figure 5A and B), suggesting that a shift in the phosphorylation state of VP30 toward hyperphosphorylation impaired EBOV propagation.

**Figure 5.**
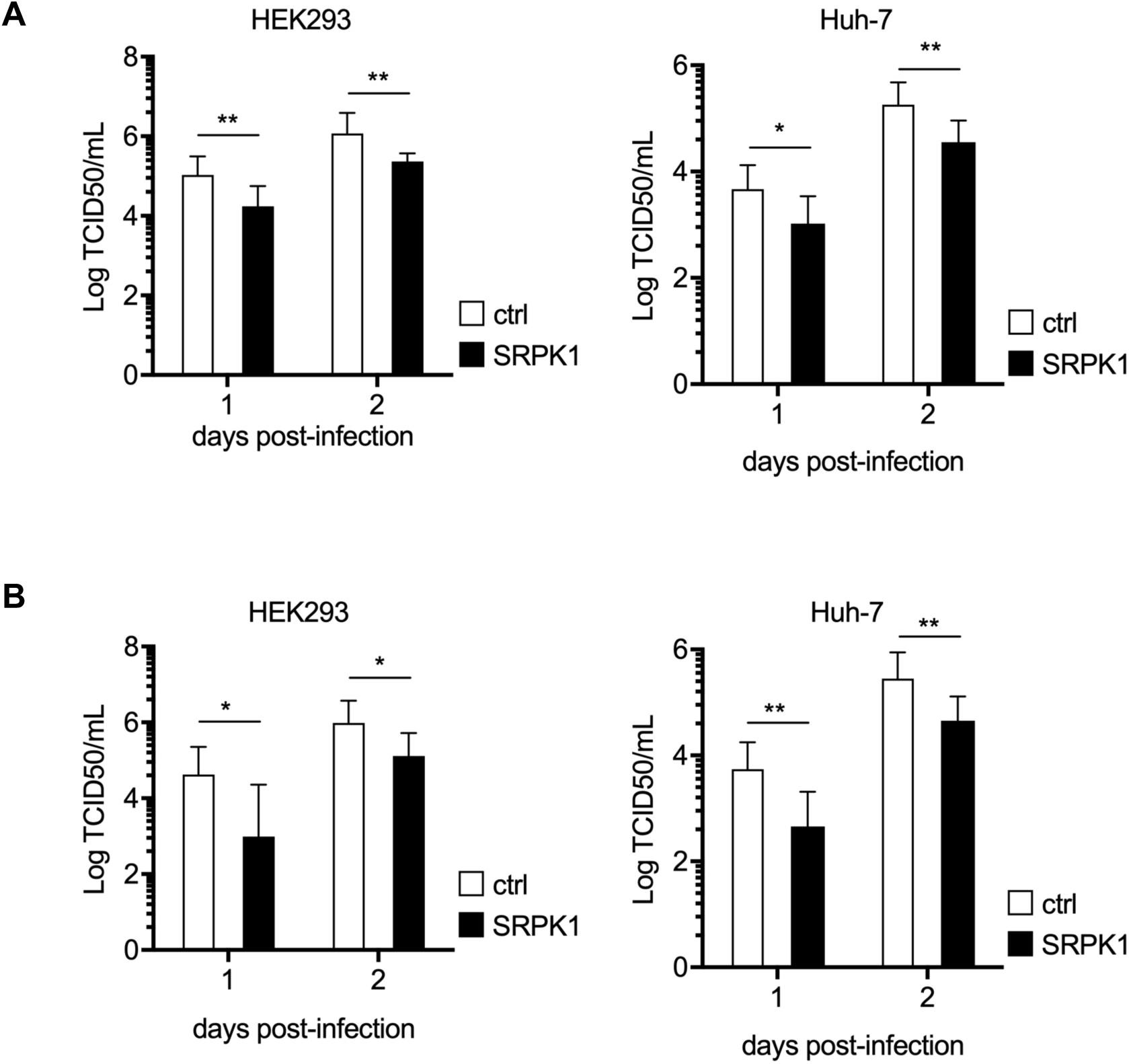
Influence of ectopic expression of SRPK1 on EBOV infection. (A, B) HEK293 or Huh-7 cells were transfected with the SRPK1-encoding plasmid 12 h prior to infection with recEBOV_wt or recEBOV_29S at an MOI of 0.1. Cell supernatants were collected at one and two days p.i. for titration of the amount of infectious virus particles released (TCID50). The mean and SD of three independent experiments carried out with recEBOV_wt (A) and recEBOV_29S (B), respectively, are displayed. Asterisks indicate statistical significance; *p < 0.05, **p < 0.01.

Next, we analyzed the effect of SRPIN340 on EBOV and found that treatment with the inhibitor did not result in significant changes in EBOV growth (Supplementary Figure 4), suggesting that, in addition to SRPK1 and SRPK2, other cellular kinases have impact on the phosphorylation state of VP30 and thus on its transcriptional support activity.

In summary, ectopic expression of SRPK1 impaired EBOV propagation, likely through hyperphosphorylation of VP30, whereas SRPIN340-mediated inhibition of endogenous SRPK1 had no significant effects on EBOV infection suggesting the presence of alternative VP30-specific kinases.

### Characterization of the interaction between VP30 and SRPK1

The recognition motif for SRPK1 has not been clearly identified. However, the motif RxxS, which is present three times in the N-proximal serine clusters of VP30, is a consensus sequence for many kinases (30, 31) (Figure 6A). Previous work has revealed that serine-29 and −31 of VP30 are the main phosphoacceptor sites that influence its transcriptional activity(19). Both serines are located in an RxxS motif. Conversely, serine-42, which is also part of an RxxS motif, does not appear to be involved in regulating transcription (19).

**Figure 6.**
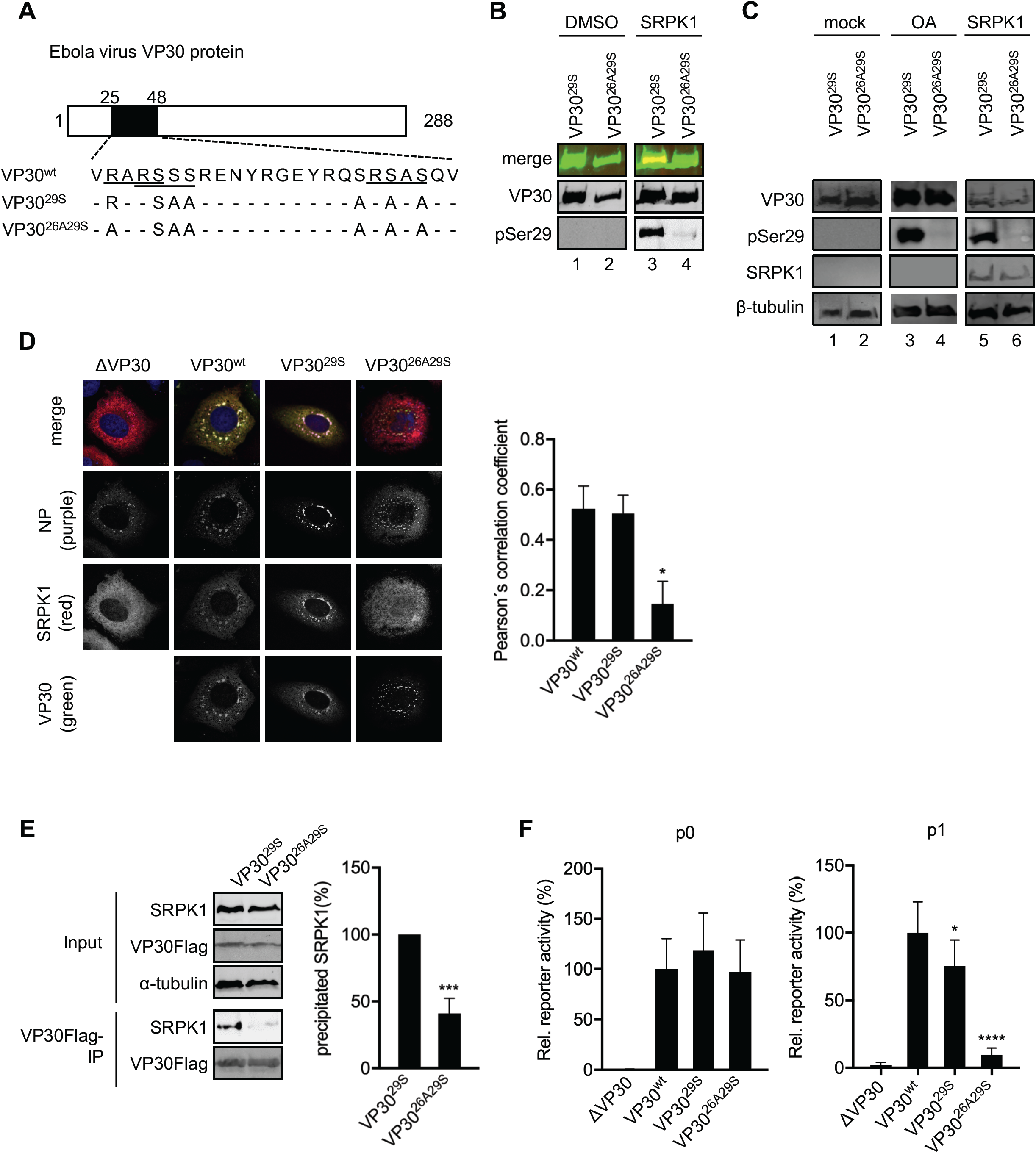
Impact of the SRPK1 binding motif R_26_xxS_29_ on the activity of VP30 in EBOV transcription and growth. (A) Schematic drawing of Ebola virus VP30 showing amino acid sequences 25-48 of VP30 and mutants of VP30. (B) *In vitro* phosphorylation of purified VP30^29S^ or VP30^26A29S^ by SRPK1 (see Figure 1B). (C) SRPK1 coexpression affects the phosphorylation status of S29. HEK293 cells were transfected with plasmids encoding VP30^wt^, VP30^29S^, or VP30^26A29S^. Cells were treated with either DMSO (control), OA, or transfected with 500 ng of a plasmid encoding SRPK1. Cell lysates were subjected to SDS-PAGE and western blotting. (D) Huh-7 cells transfected with plasmids encoding NP, SRPK1, SRPK1-TagRFP, and either VP30^wt^, VP30^29S^ or VP30^26A29S^ were fixed at 24 h p.t. and analyzed by confocal immunofluorescence microscopy. The intracellular distributions of NP, SRPK1 and VP30 were analyzed using specific antibodies and autofluorescence (SRPK1-TagRFP). Graph: Quantification of colocalization signals by Pearson’s correlation coefficients for SRPK1 and VP30. The mean and SE from 16 images derived from three independent experiments are shown. (E) HEK293 cells were transfected with plasmids encoding SRPK1 and Flag-tagged VP30^29S^ or Flag-tagged VP30^26A29S^ proteins. At 48 h p.t., the cells were lysed, and protein complexes were precipitated using mouse anti-Flag M2 agarose. An aliquot of the cell lysates (input) was collected before precipitation. Precipitates and input cell lysates were analyzed by SDS-PAGE and western blotting using SRPK1-, Flag-, and α-tubulin -specific antibodies. Graph: The amount of coprecipitated SRPK1 is indicated as %. The amount of SRPK1 coprecipitated by VP30^29S^ was set to 100%. The mean and SD from 3 independent experiments are indicated. (F) HEK293 cells were transfected with plasmids encoding trVLP assay components. Relative reporter activities are shown in p0 producer cells (analyzed at 72 h p.t.) and p1 indicator cells (analyzed at 48 h p.i.). The luciferase activity of VP30^wt^-expressing cells was set to 100%. The mean and SD from three independent experiments are indicated. Asterisks indicate statistical significance; *p < 0.05, ***p < 0.001, and ****p < 0.0001.

To investigate whether SRPK1 recognizes serine residues of VP30 within an RxxS motif, we focused on the R_26_ARS_29_ motif, including the key serine at position 29 (19, 20). A mutant of VP30^29S^ was constructed by replacing arginine-26 with alanine (VP30^26A29S^, Figure 6A), with the expectation that VP30^26A29S^ would not be phosphorylated by kinases that recognize the RxxS motif as a substrate. *In vitro* phosphorylation assays showed that recombinant SRPK1 efficiently phosphorylated VP30^29S^ but not VP30^26A29S^ (Figure 6B, lane 3 vs. 4). When VP30^29S^ or VP30^26A29S^ was ectopically expressed in HEK293 cells, neither protein was phosphorylated, indicating that endogenous SRPK1 was not sufficient to counteract the activity of PP2A, supporting the results presented in Figure 2A (Figure 6C, lanes 1, 2). However, when the phosphatases were inhibited by OA, VP30^29S^ was phosphorylated, but VP30^26A29S^ was not (Figure 6C, lane 3 vs. 4). Moreover, ectopic coexpression of SRPK1 resulted in phosphorylated VP30^29S^ but not phosphorylated VP30^26A29S^ (Figure 6C, lane 5 vs. 6). These results suggest that replacement of the arginine at position 26 by alanine destroyed the substrate sequence of SRPK1 as well as any other potential VP30-specific kinases that phosphorylate serine-29. To substantiate these findings, we analyzed the intracellular distribution of VP30 and SRPK1 by immunofluorescence and their interaction by coimmunoprecipitation. In the presence of NP, VP30 was recruited into NP-induced inclusion bodies; in contrast, expression of VP30 alone results in a homogenous cytoplasmic distribution (10). Upon coexpression of NP and SRPK1, the latter was diffusely distributed in the cytoplasm, whereas NP was detected mainly in perinuclear inclusions (Figure 6D). However, when we additionally expressed VP30^wt^ or VP30^29S^, SRPK1 was recruited into inclusion bodies together with VP30. In contrast, when VP30^wt^ was replaced by VP30^26A29S^ SRPK1 was not recruited into the NP-induced inclusions (Figure 6D). Quantification of VP30 and SRPK1 colocalization revealed a high Pearson’s correlation coefficient for VP30^wt^ and VP30^29S^ (0.52 and 0.50, respectively), which was significantly reduced for VP30^26A29S^ (0.14) (Figure 6D, right graph). These results indicate that SRPK1 was corecruited by VP30 into NP-induced inclusion bodies depending on the intact R_26_xxS_29_ motif. To confirm the interaction between SRPK1 and VP30, we performed coimmunoprecipitation analyses of ectopically expressed SRPK1 and VP30^29S^ or VP30^26A29S^ and found that VP30^26A29S^ precipitated significantly less SRPK1 compared to VP30^29S^ (Figure 6E).

Next, we analyzed the impact of a mutated R_26_xxS_29_ motif in the trVLP assay (26). While the mutated kinase recognition motif in VP30^26A29S^ had no impact on secondary transcription in producer cells (Figure 6F, producer cells), primary transcription in indicator cells was significantly reduced when the cells were infected with VLPs containing VP30^26A29S^ (Figure 6F, VP30^26A29S^, indicator cells).

These results emphasize the significance of serine-29 as a target of phosphorylation/dephosphorylation processes to regulate primary viral transcription and also identify arginine-26 as a crucial residue for serine-29 phosphorylation by kinases recognizing RxxS motifs, such as SRPK1.

### Impact of the kinase recognition motif R_26_xxS_29_ of VP30 on EBOV propagation

To analyze the influence of the R_26_xxS_29_ motif during EBOV infection, we employed three different recombinant EBOVs encoding either VP30^26A29S^, VP30^29S^ or VP30^wt^ (recEBOV_26A29S, recEBOV_29S (19), recEBOV_wt). Based on the results shown in Figure 6, recEBOV_26A29S should not be able to functionally interact with SRPK1 or other kinases that recognize the RxxS motif. In fact, compared to recEBOV_wt (circles) and recEBOV_29S (squares), viral titers in the supernatant of infected HEK293 cells were reduced by up to 2-3 log for recEBOV_26A29S (triangles, MOI of 0.1, Figure 7A). Strongly reduced growth for recEBOV_26A29S in comparison to recEBOV_wt and recEBOV_29S was also detected in Huh-7 cells (Figure 7B). These results were confirmed using a lower MOI (0.01, Supplementary Figure 5). In addition, electron microscopic analyses showed all three recombinant viruses to be morphologically indistinguishable (Figure 7C). Furthermore, colocalization analysis of VP30 and SRPK1 in cells infected with the recombinant EBOVs revealed a high Pearson’s correlation coefficient for recEBOV_wt- and recEBOV_29S-infected cells (0.48 and 0.55, respectively), but the correlation coefficient was reduced for recEBOV_26A29S-infected cells (0.18, Figure 7D). These findings indicate impaired interaction between VP30 and SRPK1 in recEBOV_26A29S infection. As we have shown that VP30^26A29S^ is active in a minigenome assay where only secondary transcription of a single gene is monitored, the observed inhibition of recEBOV_26A29S is most likely due to the interruption of primary transcription in newly infected cells and possibly by inhibiting the reinitiation of mRNA synthesis at internal genes.

**Figure 7.**
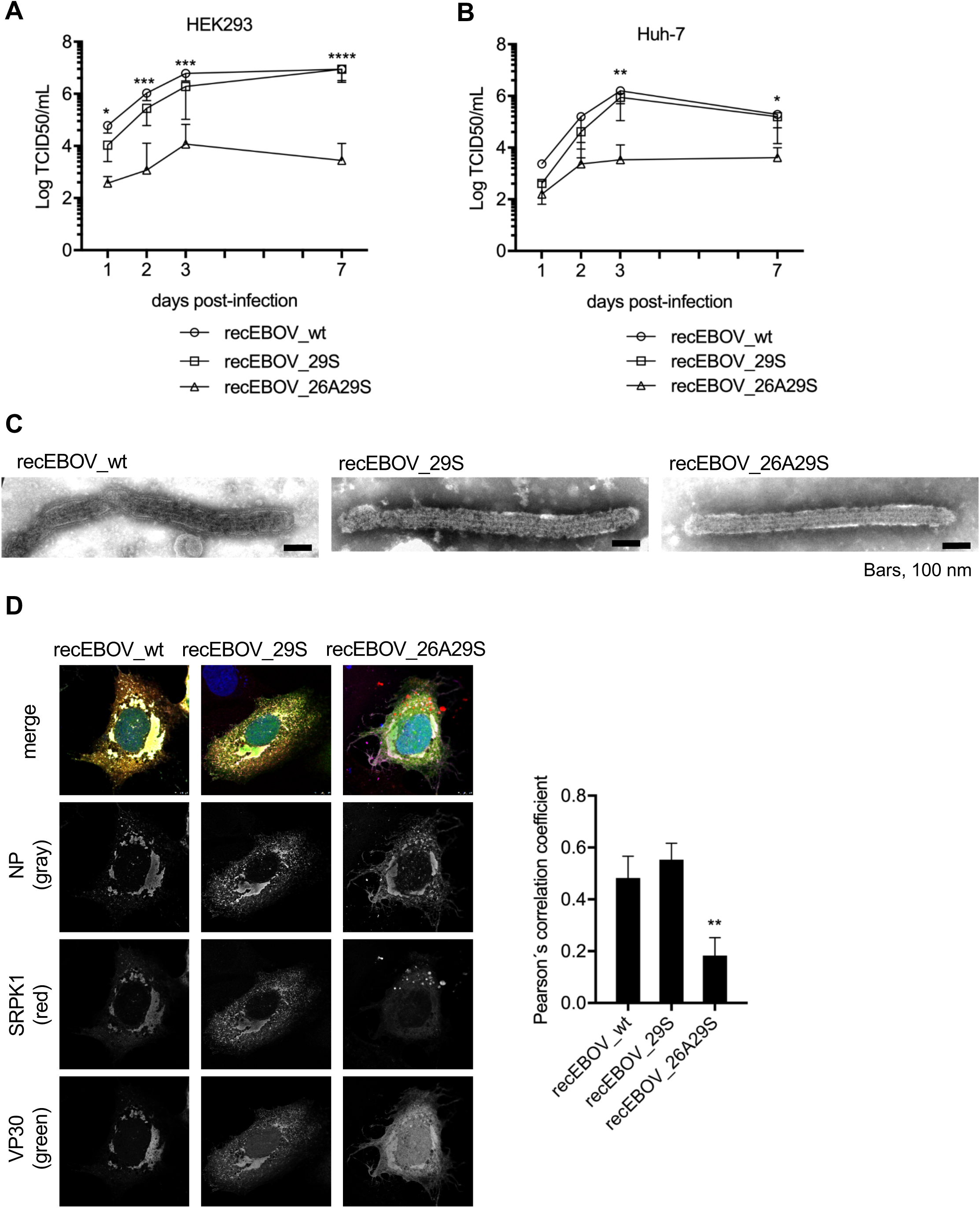
Defective replication of recEBOV_26A29S. (A, B) HEK293 (A) or Huh-7 (B)cells were infected (MOI of 0.1) either with recEBOV_wt, recEBOV_29S, or recEBOV_26A29S. The inoculum was removed at 1 h p.i., and cell supernatants were collected at 1, 2, 3, and 7 d p.i. for titration of infectious virus particles released (TCID50). The mean and SD from three independent experiments are indicated. o: recEBOV_wt, □: recEBOV_29S, Δ: recEBOV_26A29S. (C) Electron microscopy analyses of released viral particles. HEK293 cells were infected with either recEBOV_wt, recEBOV_29S, or recEBOV_26A29S. Cell supernatants were collected at 48 h p.i., and viral particles were purified via ultracentrifugation and fixed with paraformaldehyde. Samples were subjected to negative staining and analyzed by transmission electron microscopy. (D) Huh-7 cells expressing SRPK1 and SRPK1-TagRFP were infected with recEBOV_wt, recEBOV_29S, or recEBOV_26A29S. After 1 h of infection, the inoculum was removed, and the cells were fixed at 24 h p.i. for analysis of the intracellular distribution of NP, VP30 and SRPK1 by confocal immunofluorescence microscopy. Graph: Quantification of colocalization signals by Pearson’s correlation coefficients for SRPK1 and VP30. The mean and SE of 16 images derived from 3 independent experiments are shown. Asterisks indicate statistical significance; *p < 0.05, **p < 0.01, ***p < 0.001, and ****p < 0.0001.

Overall, the results indicate that SRPK1 interacted directly with VP30 and was recruited into NP-induced inclusion bodies in EBOV-infected cells. SRPK1 specifically recognized and phosphorylated serine-29 of VP30 in a manner dependent on the R_26_xxS_29_ motif, which is conserved among all Ebola viruses. SRPK1 activity regulated EBOV transcription, and mutation of the substrate motif significantly inhibited the growth of recombinant EBOVs containing serine-29 as the only phosphorylatable site at the N-terminus of VP30. Thus, SRPK1 is an important VP30-specific kinase contributing to the complex phosphorylation/dephosphorylation steps involved in EBOV transcription.

## Discussion

EBOV VP30 has been described as a viral phosphoprotein and recent studies showed that phosphorylation of VP30 need to be tightly regulated (10, 15, 16, 18). Nonphosphorylated VP30 is essential and sufficient to support secondary transcription in an EBOV-specific minigenome system and, indeed, the majority of VP30 molecules in EBOV-infected cells are nonphosphorylated (20). Importantly, however, phosphorylatable serine residues near the N-terminus of VP30 are still necessary and sufficient to support primary viral transcription as well as the reinitiation of VP30-mediated transcription at internal EBOV genes (19, 32). Serine residue 29 is most important in this respect (19). Thus, a cycle of phosphorylation and dephosphorylation events is key to support the transcriptional activity of EBOV VP30 (10, 16, 19, 32).

The present study builds on these observations, demonstrating that SRPK1 and SRPK2 were able to phosphorylate VP30 and the SRPK1/2-specific inhibitor SRPIN340 significantly reduced phosphorylation of serine-29 (Figures 1 and 2). In addition, SRPIN340 inhibited primary viral transcription in an EBOV-specific trVLP assay indicating the biological relevance of SRPK1-mediated phosphorylation of VP30 (Figure 4B, Supplementary Figure 4).

We showed that SRPK1 uses the kinase recognition motif R_26_ARS_29_ to phosphorylate serine-29 of VP30 and as a tool for its recruitment into the EBOV inclusion bodies where transcription and replication take place. The R_26_xxS_29_ motif is conserved among all ebolavirus species (Supplementary Figure 6), indicating an important role for the function of VP30. Indeed, our study demonstrated that mutation of the R_26_ARS_29_ motif in recombinant EBOV excluded SRPK1 from the inclusion bodies and dramatically impaired EBOV propagation (Figure 7A, B, D, Supplementary Figure 5).

SRPK1 and SRPK2 are cellular kinases that shuttle between the cytosol and nucleus and target proteins containing serine–arginine-rich domains (SR proteins, for a review, see Czubaty *et al*. (33)). It has been reported that SRPK1 also phosphorylates viral proteins (34-38). For example, SRPK1 is recruited by the E1^E4 proteins of human papillomaviruses and ICP27 protein of herpes simplex virus 1, resulting in downregulation of host SR protein phosphorylation (34, 36). With regard to RNA viruses, inhibition of SRPK1 by SRPIN340 suppresses hepatitis C virus and human immunodeficiency virus replication, though these effects might be caused by modulation of RNA splicing through alteration of the SR protein phosphorylation status (24, 25).

The results presented herein complement previous studies on the role of the cellular phosphatase PP2A that specifically dephosphorylates NP-associated VP30 (22). We presume a delicate interplay between PP2A and SRPK1/SRPK2 to regulate primary and secondary transcription of EBOV (22).

Figure 8 shows the current working model of how VP30 can be phosphorylated and dephosphorylated to ensure its function in EBOV transcription. Nucleocapsid-associated VP30 in released virions is phosphorylated (15). Once the virus envelope has fused with the endo/lysosomal membrane during the entry process and the nucleocapsid entered the cytoplasm, phosphorylated VP30 is dephosphorylated by PP2A, which is recruited via its subunit B56 to the LxxIxE motif of NP in close proximity to the VP30-binding motif PPxPxY (21, 22) (➀, boxed area illustrates the location of the binding motifs on NP). Dephosphorylation weakens binding between VP30 and NP, and released dephosphorylated VP30 is directed toward the first transcriptional start site (tss) of the viral RNA (tss-NP ➁) (17, 39-41). At tss-NP VP30 recruits the polymerase complex via binding of VP35 (16, 42) and initiates transcription (➂). Subsequently, VP30 is phosphorylated by SRPK1 and thereby released from its interaction with RNA and VP35 (➂, boxed area) (17). After another round of dephosphorylation by NP-associated PP2A (➃), VP30 moves to the next transcription start site (tss-VP35) to re-initiate transcription of the second gene (➄). Overall, the interplay of SRPK1 and PP2A is presumed to provide a full regulatory circuit to ensure VP30’s activity in primary EBOV transcription.

**Figure 8.**
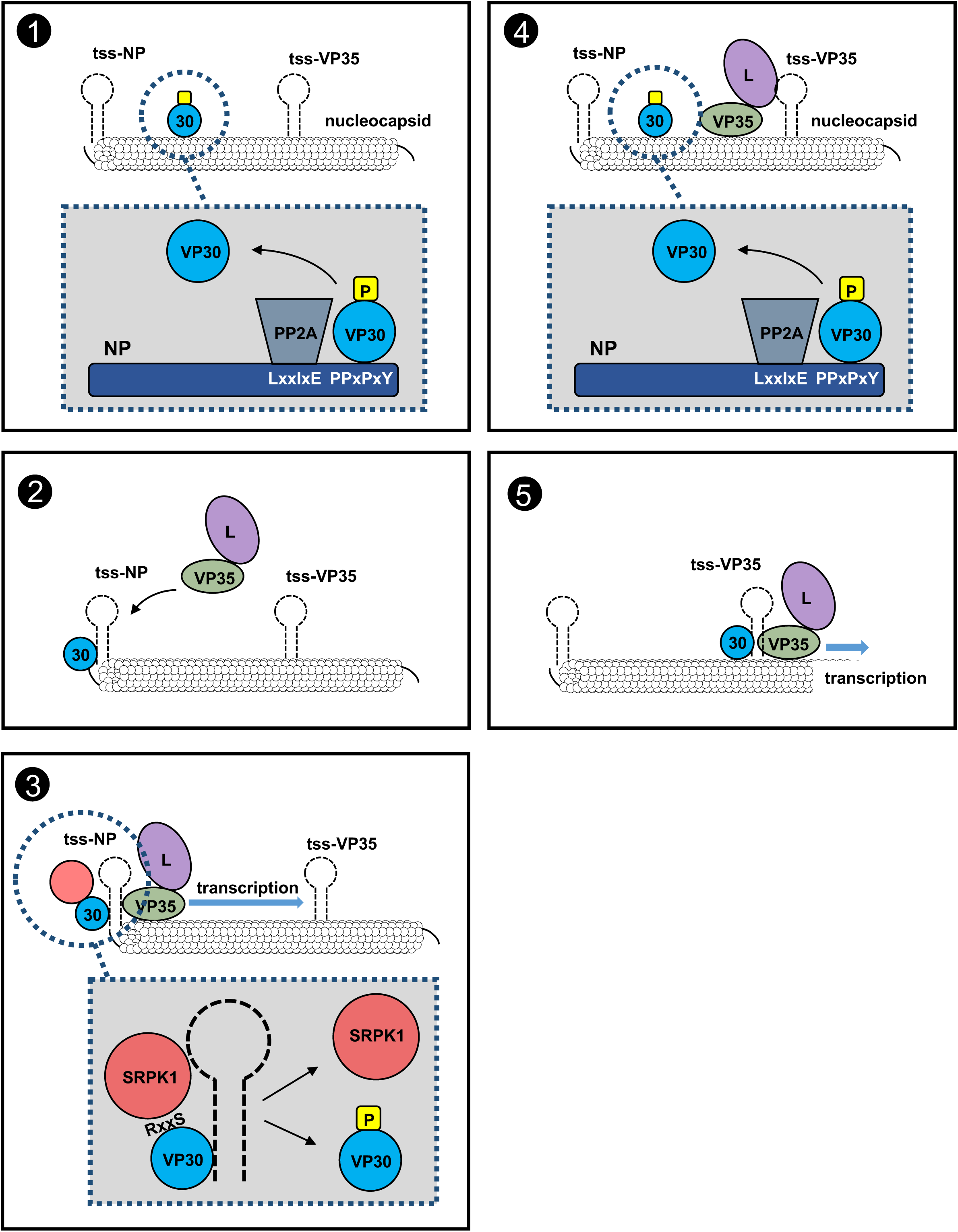
Model of VP30 reversible phosphorylation during the EBOV replication cycle. NP expression induces inclusion body formation. A sequence of phosphorylation and dephosphorylation events is essential for a fully functional VP30 in primary and secondary transcription. Within NP-induced inclusion bodies, SRPK1 is corecruited with VP30, which harbors SRPK1 recognition sites at its N-terminus site (➀). EBOV NP recruits subunit B56 of the cellular phosphatase PP2A by interaction with a motif in the direct vicinity of the VP30-binding site. VP30 is dephosphorylated (➁) and associates with the polymerase cofactor VP35 by RNA interaction and clamps the RNA template together with polymerase L and VP35 (➂). Transcription of viral RNA is initiated. The host kinase SRPK1, recruited to the viral inclusions by VP30 through interaction of the conserved RxxS motif in Ebola viruses, enables transient phosphorylation of VP30. Phosphorylated VP30 loses RNA-binding affinity and dissociates from the polymerase complex (➃), binds to NP before being again dephosphorylated by PP2A and redirected to the second transcriptional start site to reinitiate transcription of the second gene (➄). The interplay of SRPK1 and PP2A provides a full regulatory circuit to ensure VP30’s activity in EBOV transcription. Letters in the NP helix emphasize the PP2A-B56-binding motif LxxIxE and the VP30-binding motif PPxPxY. tss. transcription start site.

## Methods

### Cell culture

Huh-7 (human hepatoma), HEK293 (human embryonic kidney), VeroE6 (African green monkey kidney) cells were maintained in Dulbecco’s Modified Eagle Medium (DMEM, Life Technologies) supplemented with 10% (vol/vol) Fetal bovine serum (FBS, PAN Biotech), 5 mM L-glutamine (Q; Life Technologies), 50 U/mL penicillin, and 50 μg/mL streptomycin (PS; Life Technologies) and grown at 37 °C with 5% CO_2_.

### Plasmids

All plasmids coding for wild-type EBOV proteins (pCAGGS -NP, -VP35, -VP40, -GP, -VP30, -VP24, and -L), the EBOV-specific minigenome (3E5E-luc), pCAGGS -T7 polymerase, have been described earlier (14, 26). The plasmids of pCAGGS-VP30^29S,^ - VP30^6D^, and -VP30^6A^ were also described before (10, 19). Cloning of pCAGGS-VP30^26A29S^ was performed with the multi-site directed mutagenesis kit (Agilent) according to the manufacture’s recommendations. Construction of the pBADM41/VP30 _8-272_ variant plasmids (VP30^wt^, -VP30^6D^, -VP30^6A^, -VP30^29S^, -VP30^26A29S^) used for bacterial purification of Maltose-binding protein MBP-VP30 _8-272_ have been described earlier (17). Human SRPK1 DNA was derived from pDONR223-SRPK1, a gift from William Hahn & David Root (Addgene plasmid # 23582), and cloned into pCAGGS vector plasmids. TagRFP-SRPK1 derivatives were produced by using ligation-PCR technique as indicated previously (43). All constructs were verified by sequencing. Primer sequences are available upon request.

### Co-immunoprecipitation and Mass spectrometric analysis

Co-IP of EBOV components was carried out as previously described (16) with following modifications. For co-immunoprecipitation, HEK293 cells in 6 well plates were transfected with 500 ng/well of each protein-coding plasmid by using TransIT (Mirus). After 48 h, cells were lysed with 500 μL of ice-cold co-immunoprecipitation lysis buffer [20 mM Tris-HCl, 100 mM NaCl, 1% Nonidet P-40, 17.5 mM EDTA, and 0.1% Triton X-100, pH 7.5, with Complete protease inhibitor mixture (Roche)] for 20 min at room temperature, then subjected to sonication for 3 × 20 s at 4°C (Branson Sonifier 450). The clarified supernatant was added to 45 μL of equilibrated mouse anti-Flag M2 affinity gel agarose (Sigma-Aldrich). The reaction was incubated overnight at 4°C on a lab rotator. Elution of precipitates was achieved with 60 µL of elution buffer [125 mM Tris-HCl, 2% SDS, pH 6.8] for 5 min at room temperature, and subjected to 95 °C for 3 min. The agarose beads were completely removed by two rounds of centrifugation (14,000 rpm for 5 min at 4 °C). The final supernatant was subjected to SDS-PAGE.

Samples for mass spectrometry were prepared as follows. HEK293 cells (150 cm^2^ dish) were transfected with 6 μg of each protein-coding plasmid by using TransIT (Mirus). Cell lysis and elution were performed as described above. For loading the samples onto an SDS-PAGE, glycerol was added to the eluted supernatants and PAGE was started. Immediately after samples had entered the separation gel, electrophoresis was stopped. After staining the gel with colloidal coomassie, the protein bands were excised and subjected to in-gel digestion with trypsin (44). Mass spectrometric analysis was performed as described (45) using an Orbitrap Velos Pro mass spectrometer (Thermo Fisher Scientific) which was connected online with a nano C18 RP column equipped Ultimate nanoRSLC-HPLC (Rapid separation liquid chromatography-high performance liquid chromatography) system (Dionex). An aliquot of 15 µL of the tryptic digest was usually injected onto a C18 pre-concentration column and automated trapping and desalting of the sample was performed at a flowrate of 6 µL/min using water/0.05% formic acid as solvent.

Tryptic peptides were separated with water/0.045% formic acid and 80% acetonitrile/0.05% formic acid at a flow rate of 300 nL/min. The column was connected to a stainless steel nanoemitter (Proxeon, Denmark) and the eluent sprayed directly towards the heated capillary of the mass spectrometer using a potential of 2300 V. A survey scan with a resolution of 60,000 within the Orbitrap mass analyzer was combined with at least three data-dependent MS/MS scans with dynamic exclusion for 30 s either using CID (Collision-induced dissociation) with the linear ion-trap or using HCD (Higher energy collisional dissociation) and orbitrap detection at a resolution of 7,500. Data analysis was performed using Proteome Discoverer (v4.0; Thermo Fisher Scientific) with SEQUEST and MASCOT (v2.4; Matrix science) search engines using either SwissProt or NCBI databases (46).

### Expression and purification of maltose-binding protein fusion proteins

The procedure for expression and purification of MBP VP30 _8-272_ proteins (-VP30^wt^, - VP30^6D^, -VP30^6A^, -VP30^29S^, -VP30^26A29S^) were described earlier (17). The purity of the protein was analyzed by SDS-PAGE and Coomassie Blue staining. Protein concentration was measured by Pierce Coomassie (Bradford) Protein Assay Kit (Thermo Fisher Scientific).

### Phosphorylation detection assay *in vitro* and in cells

The *in vitro* reaction between each purified mutant of VP30 (-VP30^wt^, -VP30^6D^, -VP30^6A^, -VP30^29S^, -VP30^26A29S^) and the employed kinases was analyzed in 20 µL of kinase reaction buffer [40mM Tris-HCl, 20mM MgCl_2_, 0.1mg/mL BSA, pH7.5] with 2 mM ATP. The recombinant SRPK1 and PKR were purchased from Thermo Fisher Scientific, and RIOK2 was purchased from abcam. Okadaic acid (OA, Calbiochem) and SRPIN340 (Sigma-Aldrich) were dissolved in dimethyl sulfoxid (DMSO) according to the manufacturer’s recommendation. A phosphospecific antibody recognizing specifically VP30 phosphorylated at serine 29 (anti-pS29) was used for evaluation of VP30 phosphorylation (20).

The impact of endogenous kinases on phosphorylation of VP30 was analyzed in HEK293 cells (6-well-plate, seeded with 8 × 10^5^ cells/well) transfected with 500 ng of each plasmid encoding VP30 and mutants of VP30 (pCAGGS-VP30^wt^, -VP30^6D^, -VP30^6A^, -VP30^29S^, - VP30^26A29S^). At 48 h p.t., cells were lysed in TMT-Buffer [50 mM Tris-HCL, 5 mM MgSO_4_, 1 mM DTT, 0.5% Triton×100, pH7.5] with or without phosphatase inhibitor [25 nM of OA (Sigma-Aldrich) with 2 mM of ATP (Sigma-Aldrich)] and incubated for 1 h at 37 °C. The reaction mixture was sonicated for 3 × 20 s, and subsequently spun down at 14,000 rpm at 4 °C for 10 mins. The supernatants were subjected to Western blot analyses. The effect of ectopically expressed SRPK1 on phosphorylation of VP30 was analyzed in HEK293 cells expressing the different VP30 mutants (see above) which were co-transfected with a 500 ng of pCAGGS-SRPK1. At 48 h p.t., cells were lysed in TMT-Buffer for 20 min at room temperature, and subsequently sonicated for 3 × 20 s. After centrifugation at 14,000 g at 4 °C for 10 mins, supernatants were subjected to Western blot analyses.

### Antibodies

Immunofluorescence analysis: Chicken anti-NP polyclonal antibody (19) (1:250), guinea pig anti-VP30 antibody (16) (1:250) and a rabbit anti-VP30 antibody (16) (1:250). Corresponding secondary antibodies were goat anti-guinea pig Alexa Fluor 488 (ThermoFisher Scientific, 1:500), goat anti-rabbit Alexa Fluor 488 (ThermoFisher Scientific, 1:500), goat anti-rabbit Alexa Fluor 488 (ThermoFisher Scientific, 1:500), donkey anti-chicken-IRDye680RD (Li-COR, 1:500) was used. Western blot analysis: Chicken anti-NP polyclonal antibody (19) (1:500), a monoclonal mouse anti-VP40 antibody (47) (1:500), a guinea pig anti-VP30 antibody (16) (1:500), a rabbit anti-pSer29 antibody (20) (1:100), a rabbit anti-SRPK1 antibody (abcam, 1:250), and a mouse anti-alpha Tubulin antibody (abcam, 1:500). Corresponding secondary antibodies were: Donkey anti-chicken IRDye 680RD (Li-COR, 1:500), donkey anti-chicken IRDye 800CW (Li-COR, 1:500), donkey anti-guinea pig IRDye 800CW (Li-COR, 1:500), goat anti-rabbit IRDye 680RD (Li-COR, 1:500), donkey anti-rabbit IRDye 680RD (Li-COR, 1:500), swine polyclonal anti-rabbit immunoglobulins/HRP (DAKO, 1:500), donkey anti-rabbit IRDye 800CW (Li-COR, 1:500), donkey anti-mouse IRDye 680RD (Li-COR, 1:500), goat anti-mouse IRDye 800CW (Li-COR, 1:500).

### SDS-PAGE and Western blot analysis

SDS-PAGE and Western Blot analysis were performed as described previously (16, 48). The visualization of the signals was performed with Image Lab™ software for HRP-conjugated secondary antibodies or Li-Cor Odyssey imaging system for fluorescently labeled secondary antibodies as indicated in the antibodies section. The intensity of the obtained signals was analyzed by the Fiji software package (v.1.52e) (49).

### EBOV-specific minigenome assay

EBOV specific minigenome assay was performed as described previously (50). Briefly, the plasmids for minigenome assay (125 ng of pCAGGS-NP, 125 ng of pCAGGS-VP35, 100 ng of pCAGGS-VP30, 1000 ng of pCAGGS-L, with 250 ng of a EBOV-specific minigenome encoding the *Renilla* luciferase reporter gene (3E5E-luc), 250 ng of pCAGGS-T7 polymerase, and 100 ng of pCAGGS vector coding firefly luciferase reporter gene for normalization) were transfected in HEK293 cells seeded in 6-well plates. Reporter activity was measured at 48 h p.t. Optionally, the cells were treated either with 0.05% DMSO (control), 25 nM OA, or a mixture of 30 µM of SRPIN340 and 25 nM OA for 18 hours. At 48 h p.t., cells were lysed and reporter gene activity was monitored by luciferase assay (*pjk*, Germany).

### EBOV-specific trVLP assay

For analysis of transcription/replication activity, the EBOV transcription- and replication-competent virus-like particle (trVLP) assay was performed as described earlier (26, 51) with modifications. Briefly, HEK293 cells were seeded in a 6 well plate. Each well was transfected with plasmids encoding all EBOV structural proteins (125 ng of pCAGGS-NP, 125 ng of pCAGGS-VP35, 250 ng of pCAGGS-VP40, 250 ng of pCAGGS-GP, 100 ng of pCAGGS-VP30, 60 ng of pCAGGS-VP24, 1000 ng of pCAGGS-L), 250 ng of a EBOV-specific minigenome, 250 ng of pCAGGS-T7 polymerase and 100 ng of pCAGGS vector coding firefly luciferase reporter gene for normalization. Culture supernatants of the three wells were collected at 72 h p.t., and released trVLPs were purified via ultra-centrifugation through a 20% sucrose cushion. Optionally, an aliquot of trVLPs was treated with proteinase K digestion assay as described previously (52). The producer cells were lysed at 72 h p.t. and indicator cells were lysed at 48 h p.i., subsequently, luciferase reporter assay (*pjk*, Germany) was performed.

### Cell viability assay

To evaluate cell viability, the WST-1 Cell Proliferation Assay System (TaKaRa) was used following the manufacturer’s recommendation. HEK293 cells (8×10^3^ cells per well) or Huh-7 cells (2×10^3^ cells per well) were seeded in 96-well microplates and incubated for 24 h at 37 °C with 5% CO_2_. The medium was replaced with fresh medium with either DMSO, or various concentrations of SRPIN340 and incubated at 37 °C under 5% CO_2_. After 48 h of treatment, the medium was mixed with 10% WST-1 reagent, and absorbance was measured using an Autobio PHOmo microplate reader (measurement wavelength: 450 nm; reference wavelength: 600 nm).

### Rescue of recombinant EBOV

The recombinant EBOV (rEBOVwt) used in this study was based on EBOV Zaire (strain Mayinga; GenBank accession number AF272001). Cloning and rescue of full-length EBOV was performed as described earlier (53) with modifications. Briefly, 1000 ng of full-length EBOV cDNA (based on pAMP rg ZEBOV, kindly provided by G. Neumann) and helper plasmids (125 ng pCAGGS-NP, 125 ng pCAGGS-VP35, 1000 ng pCAGGS-L, 100 ng pCAGGS-VP30, and 250 ng pCAGGS-T7) were transfected into Huh-7 cells. The cell supernatants were transferred to fresh Huh-7 cells at 7 days p.t. Once the cells showed cytopathic effect (CPE), approximately after 7 to 10 days p.i., supernatants were collected and transferred to fresh VeroE6 cells (p2). The supernatants of P2 cells which showed CPE formation were harvested, aliquoted and stored at −80°C for use in all further experiments. In order to verify the sequence of recombinant viruses, viral RNA was extracted from supernatants using QIAamp viral RNA minikit (Qiagen) according to the manufacturer’s instruction. Reverse transcription with ensuing PCR steps was performed using a Transcriptor One-Step RT-PCR kit (Roche) with EBOV-specific primers. The resulting cDNA was sequenced. All work with infectious viruses was performed in the biosafety level 4 (BSL-4) facility at the Philipps-Universität Marburg according to national laws.

### EBOV infection

Growth kinetics of recombinant EBOV were determined in HEK293 or Huh-7 cells seeded into 12-well plate at a multiplicity of infection (MOI) of 0.1 or 0.01. The inoculum was removed after 1 h and cells were cultivated in DMEM containing 3% FCS, P/S, Q at 37°C under 5% CO_2_. To analyze the effect of ectopically expressed SRPK1, cells were transfected with either 500 ng of pCAGGS-SRPK1 or empty-vector 12 h prior to infection. To analyze the effect of SRPK1 inhibitor SRPIN340 effect, the infected cells were treated with 3% FCS DMEM with either 30 μM SRPIN or 0.05% DMSO at 1 h p.i. Aliquots of the cell supernatants were collected and stored at - 80 °C until usage.

### TCID_50_ analysis

Virus titers were determined by 50% tissue culture infectious dose (TCID50) assay in Vero E6 cells as described earlier (29).

### Immunofluorescence confocal laser scanning microscopy

Immunofluorescence analysis was performed as described previously (54, 55). The microscopic images were acquired by Leica SP5 confocal laser scanning microscope using a 63× oil objective (Leica Microsystems). Cells were grown on glass coverslips and fixed with 4% paraformaldehyde at 24 hours p.t. or p.i. Optionally, a nucleus staining was achieved by using DAPI (4’,6-diamidino-2’-phenylindole). For quantification of colocalization between VP30 and SRPK1, Pearson’s correlation coefficient was calculated by using Coloc2 plugin (https://imagej.net/Coloc_2) for Fiji (v.1.52e) (49)

### Electron microscopy analysis

We performed electron microscopy analyses of virus particles purified from the supernatant of HEK293 cells infected either with recVP30^wt^, recVP30^29S^, or recVP30^26A29S^. Virions were adsorbed on finder grids, subsequently negatively stained with 2% phosphotungstic acid. Then, these grids were analyzed by using JEM1400 transmission electron microscope.

### Statistical analysis

The presented mean values and standard deviations are derived from at least three independent experiments. Statistical analysis was performed by using SPSS16.0 or GraphPad Prism (Version 7.03). Normally distributed samples were analyzed by t-test or one-way ANOVA test, and the other samples were analyzed by nonparametric test. Statistically significant differences are indicated with asterisks (*, *P* < 0.05; **, *P* <0.01; ***, *P* < 0.001, *P* < 0.0001; ****).

## Acknowledgements

We thank Dirk Becker, Sonja Heck, Astrid Herwig, Katharina Kowalski, Cornelius Rohde for technical support. BSL-4 work would not have been possible without the supervision of Markus Eickmann, Olga Dolnik, Thomas Strecker and technical support by Michael Schmidt and Gotthard Ludwig. We would like to thank Jakob Nilsson for critical reading of the manuscript and very helpful discussions. The work was supported by overseas research fellowships of the Uehara memorial foundation and Japan Society for the Promotion of Science JSPS, JSPS Grant number 18J01631, 19K16666 (to Y.T.) and by the Deutsche Forschungsgemeinschaft DFG through the SFB 1021 projects A02 and B03 (to L.K., N.B., and Stephan Becker) and the German Center for Infection Research, Emerging Infections (to V.K., N.B., and Stephan Becker) and by AMED, Research Program on Emerging and Re-emerging Infectious Diseases, by AMED Japanese Initiative for Progress of Research on Infectious Disease for global Epidemic, by JSPS Core-to-Core Program A, the Advanced Research Networks, by Grant for Joint Research Project of the Institute of Medical Science, University of Tokyo, by Joint Usage/Research Center program of Institute for Frontier Life and Medical Sciences Kyoto University, by the Daiichi Sankyo Foundation of Life Science, and by the Takeda Science Foundation (to T.N.).

## Author contributions

YT and Stephan Becker conceived the project and designed the experiments. VK, LK, CL, TN and SB supported experiments by providing materials and technical trouble shooting. YT, VK, LK performed experiments and analyses were performed by YT, SH, NB and Stephan Becker. The manuscript was written by YT, VK, LK, SH, SB, TN, NB, and Stephan Becker

